# Reduced Somatosensory Oscillatory Dynamics and Inhibition in Moderate-to-Severe Nociceptive Pain

**DOI:** 10.64898/2026.06.25.734589

**Authors:** Mahak Virlley, Yin Xi, Natalie M. Bell, Tyrell Pruitt, Lin Guo, Sloan White, Fang F. Yu, Una E. Makris, Jason Zafereo, Amil M. Shah, Elizabeth M. Davenport, Joseph A. Maldjian, Amy L. Proskovec

**Affiliations:** Magnetoencephalography Center of Excellence, University of Texas Southwestern Medical Center (UTSW), Dallas, TX, 75390, U.S.A; Advanced Neuroscience Imaging Research Laboratory, UTSW, Dallas, TX, 75390, U.S.A; Department of Radiology, UTSW, Dallas, TX, 75390, U.S.A; Department of Neuroscience, UTSW, Dallas, TX, 75390, U.S.A; Department of Biomedical Engineering, UTSW, Dallas, TX, 75390, U.S.A; Division of Rheumatology, Autoimmunity, and Inflammation, University of California San Diego, San Diego, CA, 92093, U.S.A; Department of Physical Therapy, UTSW, Dallas, TX, 75390, U.S.A; Division of Cardiology, Department of Internal Medicine, UTSW, Dallas, TX, 75390, U.S.A

## Abstract

Nociceptive pain is the most common pain condition, and moderate-to-severe nociceptive pain substantially impacts daily functioning, constituting a significant public health burden. Despite this, most studies investigating the neural mechanisms underlying somatosensory processing and inhibition have focused on other pain conditions (e.g., neuropathic, nociplastic, or mixed pain). Thus, the extent to which neural aberrancies detected in these other populations extend to or differentiate from nociceptive pain conditions remains largely unknown. In this study, 29 individuals with moderate-to-severe nociceptive pain (MSNP) and 47 pain-free (PF) controls underwent magnetoencephalography (MEG) alongside a paired-pulse somatosensory stimulation paradigm to examine somatosensory cortical processing and functional inhibition. Pain status and intensity were determined using validated pain questionnaires, painDETECT and PROMIS-29, respectively. MEG oscillatory responses were source localized via a beamformer to the primary somatosensory cortex (S1) and time series data were extracted from the peak voxel to quantify the dynamics of somatosensory gating (SG; index of cortical inhibitory processing), oscillatory response power, and spontaneous power. We found that adults with MSNP exhibit aberrant theta SG in contralateral S1 compared to PF controls, reflecting reduced functional inhibition of innocuous stimulus processing in this region. Additionally, individuals with MSNP demonstrated exaggerated gamma responses but blunted alpha responses in contralateral S1 to innocuous stimulation. Finally, individuals with MSNP were characterized by weaker spontaneous alpha in contralateral S1 that scaled with self-reported pain intensity. Together, these findings suggest that experiencing MSNP is associated with disrupted somatosensory and cortical inhibitory processing.

## 1 Introduction

Nociceptive pain experiences are the most common form of pain reported (e.g., arthritis, surgical pain, mechanical lower back pain), with moderate-to-severe nociceptive pain (MSNP) conditions providing an increased burden on daily functioning and activities (Bonezzi et al., 2020; Cao et al., 2024; Dagnino & Campos, 2022; Makris et al., 2017; Ritchie et al., 2023). Nociception describes the signal transduction cascade involved in pain, and nociceptive pain experiences, such as MSNP, refer to the unpleasant sensory and emotional experiences associated with actual or potential tissue damage (Raja et al., 2020). Although nociceptive pain conditions are highly prevalent, the central neural mechanisms supporting nociceptive pain, particularly at moderate-to-severe levels, have been comparatively less characterized than those associated with neuropathic pain (i.e., pain due to nerve damage) and nociplastic pain (i.e., pain in the absence of tissue or nerve damage) (Zebhauser et al., 2023). In addition, many prior pain studies have examined mixed pain (e.g., chronic lower back pain) samples without clearly distinguishing nociceptive, neuropathic, or nociplastic features through clinically validated pain questionnaires (e.g., painDETECT), limiting mechanistic specificity (Ahn et al., 2019; Bittencourt et al., 2022; Bonezzi et al., 2020; Day et al., 2020; Galvez-Sanchez & Reyes Del Paso, 2020; Kenefati et al., 2023; Nakawaki et al., 2018; Zebhauser et al., 2023). Importantly, individuals with nociceptive pain can develop central sensitization, in which pain may persist despite resolution of peripheral tissue damage, potentially evolving towards and embodying pain characteristics of nociplastic pain phenotypes (e.g., fibromyalgia) (Ohashi et al., 2023; Vervullens et al., 2024; Woolf, 2011). Identifying neural markers of MSNP may therefore help guide early interventions aimed at pain modulation, including psychosocial treatments or targeted physical activity interventions, and ultimately reduce long-term health burden (Day et al., 2020; McGreevy et al., 2011; Nanda, 2025; Volcheck et al., 2023).

In the brain, the initial cortical region to process tactile information, including nociception, is the primary somatosensory cortex (S1), which is implicated in both bottom-up and top-down processes following the presentation of both noxious and innocuous (i.e., non-noxious) stimuli through neural oscillatory activity (Apkarian et al., 2005; Bushnell et al., 1999; Forss et al., 1994; Gross et al., 2007; Huttunen et al., 2008; Meehan et al., 2008; Raju & Tadi, 2024; Rossiter et al., 2013; Wiesman et al., 2017; Yamada et al., 2004). Electroencephalography (EEG) and magnetoencephalography (MEG) studies have demonstrated that innocuous and noxious stimulations induce a robust and early increase in theta (∼4-8 Hz) and gamma (30+ Hz) oscillations in S1, which are thought to reflect bottom-up processing (Apkarian et al., 2005; Garcia-Larrea et al., 2003; Gross et al., 2007; Iwamoto et al., 2021; Mouraux et al., 2003; Rossiter et al., 2013; Zhang et al., 2012). Gamma oscillations typically emerge early and transiently, relating to fast, perceptual sensory processing (Gross et al., 2007; Ploner et al., 2017), whereas theta oscillatory activity tends to be sustained and linked with ongoing, integrative cognitive processes that evolve from basic sensory input (Nyhus & Curran, 2010; Zebhauser et al., 2023). On the other hand, innocuous and noxious stimulation are associated with decreases in late and sustained alpha (∼8-13 Hz) and beta (∼15-30 Hz) oscillations in S1, thought to reflect attention and prediction error, i.e., top-down mechanisms during somatosensory processing (Apkarian et al., 2005; Bardouille et al., 2010; Dockstader et al., 2010; Fries, 2015; Palmer et al., 2019; Ploner et al., 2006; Ploner et al., 2017; Wiesman & Wilson, 2020).

Interestingly, in nociplastic and mixed pain conditions, theta and gamma oscillatory dynamics serving innocuous somatosensory processing are heightened, reflecting central sensitization to pain (Kenefati et al., 2023; Zebhauser et al., 2023; Zhou et al., 2018). These changes are thought to reflect disruptions in cortical disinhibition and thalamocortical rhythms (Lim et al., 2015; McGreevy et al., 2011; Volcheck et al., 2023; Woolf, 2011). However, whether individuals with MSNP also exhibit exaggerated theta or gamma activity to innocuous sensory processing in S1 has not been investigated. Additional studies using noxious stimulation during EEG and subjective pain intensity ratings have suggested that gamma engagement in S1 is correlated with pain perception in healthy controls, but this finding was not observed in a mixed pain group (i.e., individuals with migraines) (Bassez et al., 2020). Conversely, alpha oscillations were found to be correlated with pain perception in a mixed pain group (i.e., chronic lower back pain), and increasing alpha engagement was found to correspond to subjective pain relief (Ahn et al., 2019; Babiloni et al., 2006). These findings suggest that alpha and gamma oscillations differentially encode pain and cognition in healthy adults compared to adults with various pain conditions. Additionally, specifically looking at individuals with nociceptive pain, as opposed to pain conditions that contain a mix of nociceptive and neuropathic components, can help elucidate biomarkers for pain perception (Bonezzi et al., 2020). Identifying early cortical disinhibition in MSNP can also inform studies with nociplastic pain, and signal which biomarkers are specific to MSNP (Apkarian et al., 2005; Nanda, 2025).

Beyond evaluating basic somatosensory processing, characterizing higher-order inhibitory mechanisms within the somatosensory system is likely to be clinically useful. When innocuous stimuli are delivered in pairs, somatosensory gating (SG) is observed in S1 (Cheng et al., 2016; Cromwell et al., 2008; Spooner, Eastman, et al., 2020; Spooner et al., 2019; Wiesman et al., 2017). SG is a neurophysiological comparative phenomenon in which neural responses are dynamically reduced to repeated, identical stimuli and is thought to be mediated by local feedforward inhibitory mechanisms within cortical circuitry (Cromwell et al., 2008; Smucny et al., 2015). SG is commonly assessed using paired-pulse paradigms, in which two identical innocuous stimuli are delivered in rapid succession (e.g., 500 ms apart), and the neural response magnitude following the second stimulus (Stim2) is compared to that following the first stimulus (Stim1) (Cromwell et al., 2008; Spooner, Eastman, et al., 2020). Such gating reflects an inhibitory filtering mechanism in which the brain attenuates processing of redundant sensory information to preserve neural resources for behaviorally relevant computations (Cromwell et al., 2008).

Multiple studies have assessed SG of time-domain neural evoked responses in nociplastic and complex (e.g., complex regional pain syndrome) pain conditions, linking decreased functional inhibition of evoked components via SG to increased perceived pain (Lenz et al., 2011; Lim et al., 2015; Montoya et al., 2006). Although investigations on time-domain SG provide insight into intracortical inhibition, oscillatory dynamics can provide additional insight into cognition, like attention (Wiesman & Wilson, 2020). For example, theta oscillations exhibit SG in S1, with exaggerated inhibition of Stim2 observed when attentional resources are explicitly directed toward the somatosensory stimuli, as opposed to when attention is diverted to a competing visual task (Wiesman & Wilson, 2020). Interestingly, time-domain SG does not exhibit this exaggeration during attentional engagement (Wiesman & Wilson, 2020). Further, gamma SG is also observed and highly studied in S1 as a bottom-up sensory mechanism, with attention implications in clinical populations (Proskovec et al., 2020; Spooner et al., 2019; Wiesman et al., 2021). Thus, studying SG of oscillatory responses in MSNP might provide insights into the pathophysiology of nociceptive pain conditions from a bottom-up and top-down approach.

Despite a growing body of literature characterizing alterations in oscillatory dynamics underlying SG and higher-order inhibitory functions (Arif et al., 2021; Casagrande et al., 2022; Heinrichs-Graham et al., 2023; Liu et al., 2018; Spooner et al., 2021; Spooner, Wiesman, et al., 2020; Spooner et al., 2019; Wiesman et al., 2017; Wiesman et al., 2021; Wiesman & Wilson, 2020), there are currently no similar investigations in the context of MSNP. This gap is particularly striking given that these mechanisms play a foundational role in sensory filtering and attentional modulation, which are processes central to the experience of pain (Bolton & Staines, 2012; Commodari & Guarnera, 2008; Mesulam, 1998). Herein, we begin to fill this gap by using MEG and a paired-pulse somatosensory stimulation paradigm to examine oscillatory dynamics and SG in individuals with MSNP. Specifically, we investigated oscillatory responses and SG within contralateral S1 following innocuous somatosensory stimulation. Accordingly, we hypothesized that individuals who report MSNP will exhibit aberrant gating in S1, reflecting dysfunctional bottom-up inhibitory processing. We further predicted that responses to stimulation will be characterized by exaggerated high-frequency oscillations and reduced low-frequency oscillatory activity relative to pain-free controls, based on studies implicating decreased alpha engagement in pain conditions (Ahn et al., 2019) and increased gamma responses to painful stimuli (Gross et al., 2007; Hauck et al., 2007; Zhang et al., 2012). Based on previous studies implicating SG and alpha with pain perception (Ahn et al., 2019; Babiloni et al., 2006; Lim et al., 2015), we predict that alpha oscillations and SG will correlate with perceived ongoing pain intensity.

## 2 Methods

### 2.1 Study, ethics, and participants

The present analyses draw on data from the Dallas Hearts and Minds Study (DHMS), a continuation of the Dallas Hearts Study (DHS), which is a large, longitudinal, population-based study designed to examine cardiovascular and at the most recent timepoint, neurocognitive health, with intentional enrichment of racially diverse participants (Bell et al., 2026). Since brain health metrics were only included in the most recent wave of DHMS data collection, this study design is cross-sectional. All study procedures were approved by the Institutional Review Board at the University of Texas Southwestern Medical Center, and written informed consent was obtained from all participants prior to participation.

Data relevant to the current study included demographic information, structural MRI, medical history questionnaires, pain questionnaires, and a comprehensive neurocognitive assessment battery. Individuals with and without MSNP were identified using a battery of self-report pain questionnaires (i.e., PROMIS-29, painDETECT, ACR Assessments of Fibromyalgia). Individuals were also assessed for mental health conditions, a common pain comorbidity (de Heer et al., 2014; Gerrits et al., 2012). Depression scores were indexed by Quick Inventory of Depression Symptomatology (QIDS), which is a highly reliable and valid instrument for assessing depression severity (Liu et al., 2021; Reilly et al., 2015; Rush et al., 2003). Anxiety scores were indexed by the Generalized Anxiety Disorder-7 questionnaire, which has been validated in a general population (Lowe et al., 2008; Spitzer et al., 2006). Medical history forms categorized individuals based on the exclusionary criteria.

MEG data were available for a subset of participants. This subsample reflects both logistical and technical constraints, including limited scanner availability (i.e., morning sessions were prioritized for clinical epilepsy evaluations) and standard MEG exclusion criteria. Specifically, individuals with ferromagnetic implants (e.g., cochlear implants, magnetic shunts) or non-removable metal that produced substantial signal artifact (e.g., certain dental hardware) were not eligible for MEG acquisition.

Additional exclusionary criteria across all participants included factors impacting cognition, including neurological or current neuropsychiatric disorder (e.g., major depressive disorder, generalized anxiety disorder), current high intensity alcohol use (Hingson et al., 2017), concussion history in the past year, or any medical illness affecting the central nervous system (except fibromyalgia for MSNP group only, explained below).

#### 2.1.1 Group definitions

Pain intensity was defined by the NIH pain intensity item from the PROMIS-29 global assessment: “In the past 7 days, how would you rate your pain on average?” (Hays et al., 2018). Scores ranged from 0-10. Ongoing moderate-to-severe pain was defined as a score between 4-10 on the PROMIS-29 pain intensity item. The PROMIS-29 instrument is highly reliable and valid in the assessment of health-related quality of life (DiRenzo et al., 2023; Hwang et al., 2024). Nociceptive pain was defined as a score between 0-12 on the painDETECT instrument, which is highly reliable and valid for assessing neuropathic pain (Bittencourt et al., 2022; Tampin et al., 2017). Participants in the MSNP group also reported their current pain intensity on the day of the MEG scan using the painDETECT item, “How would you assess your pain now, at this moment?”, rated on a scale from 0 (no pain) to 10 (worst imaginable pain). Some MSNP participants also endorsed mixed pain with nociplastic and nociceptive components, where the nociplastic component (i.e., fibromyalgia) was defined by the American College of Rheumatology (ACR) 2016 criteria assessed with the Patient Self-Report Survey for the Assessment of Fibromyalgia (Galvez-Sanchez & Reyes Del Paso, 2020). The ACR diagnostic criteria was found to be reliable and hold validity in differentiating between nociplastic (i.e., fibromyalgia) and nociceptive (i.e., arthritis) pain (Ablin & Wolfe, 2017; Galvez-Sanchez et al., 2020; Wolfe et al., 2016). In addition, medical history forms included a question asking if the participant was diagnosed with fibromyalgia by a health care physician. Thus, the MSNP group included individuals with strictly nociceptive pain, as well as individuals with mixed nociplastic and nociceptive pain, which was controlled for in the statistical analyses (see below).

Pain-free (PF) controls were obtained from DHMS. Pain-free category was defined as a score of 0 on the PROMIS-29 pain intensity item, a score of 0 on the painDETECT instrument, and not qualifying for the ACR diagnostic criteria based on the Patient Self-Report Survey for the Assessment of Fibromyalgia. Therefore, they did not endorse any type of pain.

### 2.2 Experimental paradigm

During MEG, participants were seated at a 60° position in a nonmagnetic chair with an accompanying desk in front of them. The MEG chair was adjusted such that their head was comfortably touching the top of the MEG helmet sensor array. Additionally, foam padding was placed around the head to minimize head motion. During recording, they were asked to keep their eyes closed and to keep their right hand and arm relaxed throughout the experiment. Electrical stimulation was applied unilaterally to the right median nerve using external cutaneous stimulators connected to a Digitimer DS7A constant-current stimulator system. One-hundred paired-pulse trials (i.e., total of 200 pulses) were collected with an inter-stimulus interval of 500 ms and inter-pair interval of 3950 ± 150 ms, resulting in a total runtime of approximately 7 minutes. Each pulse generated a 0.2 ms constant-current square wave that was set to 10% above the motor threshold required to elicit a subtle thumb twitch. Participant-specific stimulation levels were recorded and evaluated as a potential covariate in regression-based analyses, with results reported only when a significant covariate relationship was identified.

### 2.3 MEG data acquisition

The MEG and MRI data acquisition are detailed in our related protocol manuscript (Bell et al., 2026). MEG recordings were conducted in a 3-layer magnetically-shielded room using a 306-sensor MEGIN TRIUX Neo system at a sampling rate of 1 kHz and acquisition bandwidth of 0.1-330 Hz. The TRIUX Neo system contains 102 magnetometers and 204 planar gradiometers. Participants were monitored for safety during acquisition using a real-time audio and video system. Prior to recording, a 3D digitizer was used to create a map of the subject’s head including the location of nasion and preauricular fiducials and five head position indicator (HPI) coils. Additionally, during the MEG recording, an electric current with a unique frequency (e.g., 320 Hz) was fed to each HPI coil, allowing each coil to be continuously localized with reference to the sensors, and head motion to be corrected for offline. Following acquisition, MEG data was individually corrected for head motion and subjected to noise reduction using a signal space separation method with a temporal extension (Medvedovsky et al., 2007; Taulu et al., 2004; Taulu & Simola, 2006).

### 2.4 MRI acquisition, processing, and MEG-MRI coregistration

Following the MEG scan, fiducial markers that appear hyperintense on T1-weighted MRI were placed at the nasion and bilateral preauricular areas marked during the digitization step. These markers remained in place until the completion of the MRI to aid in co-registration. All participants underwent MRI using a 3 Tesla Siemens Prisma scanner with a 64-channel head coil. A whole-brain 3D T1-weighted magnetization-prepared rapid gradient echo (MP-RAGE) sequence with 1 mm^3^ isotropic resolution was acquired (repetition time, 1800 ms; echo time, 2.26 ms; field of view, 256 mm; matrix, 256 x 256; slice thickness, 1mm; acquisition plane, sagittal; flip angle, 8 degrees). Structural MRI data were processed using the FreeSurfer (version 7.1.1) recon-all pipeline to obtain segmented anatomical images. These outputs were subsequently imported into Brainstorm (version 3.4; (Tadel et al., 2011)) and normalized to standard MNI space (Fonov, 2009). Since MEG HPI coil locations were also known in head coordinates, all MEG measurements could be transformed into a common coordinate system. With this coordinate framework, each participant’s MEG data were co-registered with their T1-weighted MRI prior to source-level analysis. Source reconstruction was then performed in this aligned space, after which the resulting functional maps were projected into MNI space using the transformation parameters derived from the structural MRI normalization. Structural (i.e., surface-based morphometry) and diffusion (i.e., diffusion tensor imaging and neurite orientation dispersion and density imaging) MRI analyses were also conducted to examine potential associations with SG; detailed methods are in Supplementary Material.

### 2.5 MEG preprocessing

MEG data preprocessing, time-frequency analysis, and beamforming is detailed in our DHMS MEG protocol manuscript (Bell et al., 2026). Briefly, MEG data were notch-filtered (i.e., at 60 Hz and its harmonics) and band-pass filtered between 1 and 200 Hz. Artifacts (e.g., cardiac, ocular) were removed using independent component analysis. MEG data were visually inspected for any further contamination (e.g., muscle), and those time segments with remaining artifact were not utilized in subsequent analyses. Cleaned, continuous time-series data were split into epochs assigned as 4300 ms in length with 0 ms denoting delivery of the first stimulus and 500 ms denoting delivery of the second stimulus. Baseline was defined as -700 to -300 ms. On average 90 (SD = 7) remaining usable trials were used for the MSNP group, and 88 (SD = 7) trials were used for the PF controls in the final analyses. Trial counts did not significantly vary by group (Mann-Whitney U test, *p* > 0.05). Only data from planar gradiometers were used during analysis.

### 2.6 MEG sensor-level analysis

Artifact-free epochs were transformed into the time-frequency domain using Morlet wavelet analysis (range: 2-152 Hz, step: 1 Hz). The resulting spectrograms were averaged across trials to generate time-frequency plots of mean spectral magnitude and baseline-normalized per sensor. The time-frequency windows utilized during source reconstruction were determined by statistical analysis of the relative sensor-level spectrograms across all participants using parametric tests against a null hypothesis that the mean is zero. Spectrogram thresholds were set at *p* < 0.05, and multiple comparisons corrections using the Benjamini-Hochberg false discovery rate (FDR) procedure were applied. Only time frequency windows that contained significant oscillatory responses (i.e., differences from baseline) after FDR correction were subjected to source reconstruction, or beamforming.

### 2.7 MEG source-level analysis

Cortical source activity was reconstructed using a linearly constrained minimum variance vector beamformer, which calculates source power through spatial filtering in the time-frequency domain (Gross et al., 2001; Van Veen et al., 1997). For each participant, images were baseline-normalized using a separately averaged baseline of equal duration and bandwidth (Van Veen et al., 1997). The resulting whole-brain images, referred to as pseudo-*t* maps, contained voxel-wise pseudo-*t* values representing noise-normalized relative power at a spatial resolution of 4.0 × 4.0 × 4.0 mm. All maps were subsequently transformed into standardized MNI space and spatially resampled to 1 mm^3^ resolution. MEG preprocessing and source imaging were performed in Brainstorm (version 3.4) (Tadel et al., 2011).

The resulting voxel-wise maps of neural oscillatory response amplitude were averaged across both stimulations and all participants to generate grand-averaged images, for each frequency-specific response. From each grand-averaged map the peak voxel of maximum amplitude was identified. For each participant, a voxel-specific time series (virtual sensor) was extracted at the peak location by applying the forward model’s sensor weighting matrices to the MEG data. Relative peak voxel time series were used to derive maximum response amplitudes for the first (Stim1) and second (Stim2) stimuli, as well as the gating ratio (GR; Stim2/Stim1). Absolute peak voxel time series were additionally used to estimate spontaneous activity, defined as the mean signal magnitude during the baseline period. To prevent negative relative GR values and division by values approaching zero, frequency-specific offsets were applied to the relative stimulus amplitudes prior to GR calculation. For frequency bands exhibiting event-related synchronizations (theta and gamma), constant values were added to the relative stimulus amplitudes (35 for theta; 10 for gamma) to correct for negative values. Conversely, for bands characterized by event-related desynchronizations (ERD) (alpha and beta), constant values were subtracted (23 for alpha; 10 for beta) to offset positive values. GR was then calculated using these adjusted relative stimulus amplitudes and served as an index of SG. GR was selected over gating difference (GD; Stim1 − Stim2), as GD is insensitive to amplitude scaling effects, and these amplitude effects were hypothesized to be present in individuals with MSNP (Apkarian et al., 2005; Dagnino & Campos, 2022; Farrell, 2012; Hauck et al., 2007; Kenefati et al., 2023; Li et al., 2016; Ploner et al., 2017).

### 2.8 Statistical analyses

To probe for potential confounding variables between groups (MSNP, PF), group differences in demographic and medical history variables were evaluated. Specifically, continuous variables were compared between groups using Welch’s two-sample t-tests, which do not assume equal variances and are robust for unequal sample sizes. Categorical variables were compared using chi-square tests or Fisher’s exact tests when expected cell counts were less than 5. When assessing potential confounding variables, groups were found to significantly differ on depression scores (QIDS; *p* < 0.01) and fibromyalgia diagnosis (*p* < 0.05; Table 1). QIDS was assessed for contributing variance in all statistical analyses. That is, statistical models were run with and without its inclusion as a nuisance variable. Results were virtually identical regardless of whether QIDS was included or not and thus, it was excluded from final analyses to retain power. Fibromyalgia diagnosis was included in final analyses as a nuisance variable to control for any mixed (i.e., nociplastic) pain effects. Additionally, since age has been shown to impact pain perception (Farrell, 2012; Mullins et al., 2022; Ritchie et al., 2023), it was included as a nuisance variable in main effects models and as a variable of interest in group-by-age interactions models. Additionally, current pain intensity was assessed to evaluate whether momentary pain state contributed variance to the observed neural responses. Follow-up analyses were conducted to determine whether current pain intensity influenced the statistical outcomes. Results from these analyses are reported only when inclusion of this variable altered statistical significance.

**Table 1.**
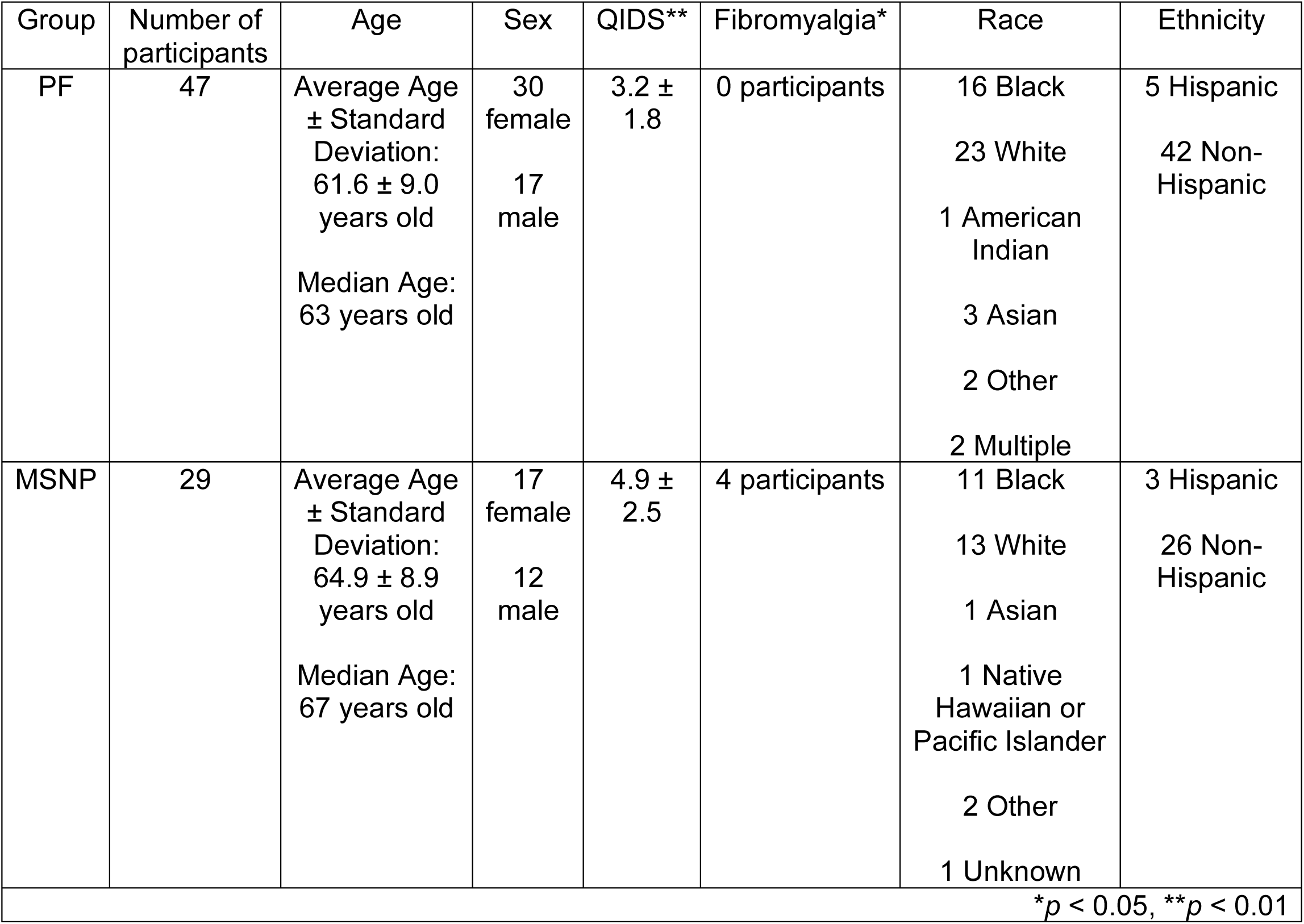
Group demographics and confounding variables (QIDS score and Fibromyalgia).

Statistical analyses focused on evaluating group differences in MEG oscillatory metrics between individuals with MSNP and those who were PF. General linear models (GLM) were used to assess group effects on MEG responses while adjusting for relevant covariates. Age and fibromyalgia diagnosis (indicated on medical history form or using the ACR diagnostic) were included as covariates in all models. For MEG metrics hypothesized to modulate ongoing pain intensity (i.e., alpha metrics and SG) (Ahn et al., 2019; Lim et al., 2015), pain measures (i.e., PROMIS-29) were incorporated as dependent variables in GLMs to test MEG oscillatory metric contributions to pain intensity, with age and fibromyalgia diagnoses controlled for, and were corrected for multiple comparisons using Bonferroni (*p* = 0.05/3 = 0.0167). Type III analysis of variance (i.e., Type III sums of squares) was applied to all models to estimate covariate-adjusted effects of group and age. Interaction terms (e.g., group × age) were examined to determine whether age moderated group differences while controlling for fibromyalgia diagnoses as a nuisance variable. Importantly, main group effects were only evaluated in absence of the interaction term. All statistical analyses were completed in R using RStudio (Posit, 2025).

## 3 Results

In DHMS, 44 individuals met criteria for MSNP and had concurrent MEG with the SG paradigm. Of these individuals, 15 were excluded due to exclusionary criteria. Thus, the final analytic sample included 29 participants with MSNP (Table 1). Within this group, 3 individuals also endorsed mixed pain (i.e., a nociplastic component) per ACR 2016 criteria (Galvez-Sanchez & Reyes Del Paso, 2020). One additional participant with MSNP indicated this diagnosis and was treated the same as those with a confirmed ACR diagnosis.

Data from 66 individuals classified as PF controls were initially included. Of the individuals, 15 were excluded due to exclusionary criteria. Four were deferred due to low English language ability. Thus, the final analytic sample included 47 PF control participants (Table 1). PF controls in the current study were a random sample of pain-free DHMS participants at the time of the analysis.

### 3.1 Sensor-level and beamformer results

Robust responses in the time-frequency domain were observed after each somatosensory stimulation in multiple oscillatory bands. We observed significant theta (4-7 Hz; 0-250 ms for Stim1, 500-750 ms for Stim2), alpha (8-13 Hz; 225-400 ms for Stim1, 725-900 ms for Stim2), beta (15-25 Hz; 125-350 ms for Stim1, 625-850 ms for Stim2), and gamma (30-90 Hz; 10-60 ms for Stim1, 510-560 ms for Stim2) responses (*p* < 0.05 corrected; Figure 1a). The responses following Stim1 and Stim2 in each band all localized to the hand-knob region of contralateral S1 (Figure 1b). Main group effects were observed in the theta, alpha, and gamma metrics, as discussed below. No significant group effects were observed regarding beta metrics.

**Figure 1.**
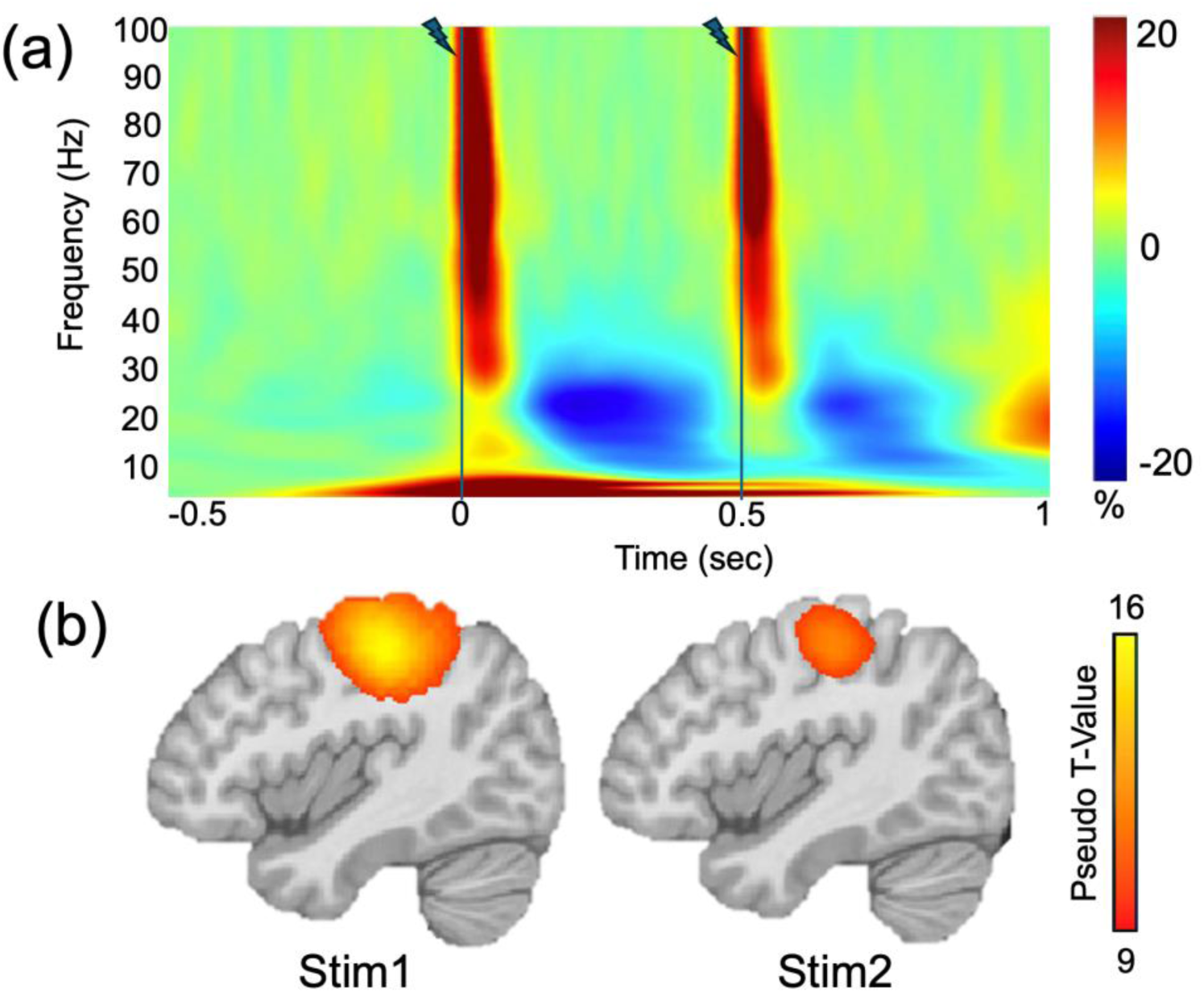
Neural Oscillatory responses to SG paradigm. (a) A grand-averaged time-frequency spectrogram derived from a MEG sensor located near the contralateral sensorimotor cortex. The x-axis indicates time in seconds. The y-axis denotes frequency in Hertz. The magnitude is shown as a percentage change unit relative to baseline with a color bar to the far right of the spectrogram. Stimulation was delivered at 0 sec and 0.5 sec, both denoted by a dark blue line and lightning bolt. (b) Gamma (30-90 Hz) beamformer images representing pseudo t-values averaged across both groups are shown for Stim1 (10-60 ms) and Stim2 (510-560 ms). Increases in gamma relative to baseline were observed in virtually identical regions in contralateral S1 following each stimulation, with the response to the second stimulation being visibly attenuated, reflecting SG.

### 3.2 Reduced theta SG in individuals with MSNP

Since SG is a functional inhibitory mechanism and pain conditions are thought to impact inhibitory processing, we evaluated whether individuals with MSNP displayed aberrant SG. Theta metrics (Stim1 relative amplitude, Stim2 relative amplitude, GR, and baseline absolute amplitude) were calculated from the peak voxel time series and entered into GLMs with group (MSNP, PF) as the variable of interest, and age and fibromyalgia diagnosis controlled for. Our results revealed a significant group effect on contralateral S1 theta SG above and beyond the effects of age and fibromyalgia (β = 0.301, t(3, 72) = 2.682, *p* = 0.009), such that individuals with MSNP exhibited less theta SG in this region relative to PF individuals (Figure 2b). The relative time series is plotted by group for visualization purposes in Figure 2c. No other theta metrics were found to significantly differ by group (all *p*’s > 0.05). Additionally, theta analyses did not yield significant group-by-age effects (all *p*’s > 0.05).

**Figure 2.**
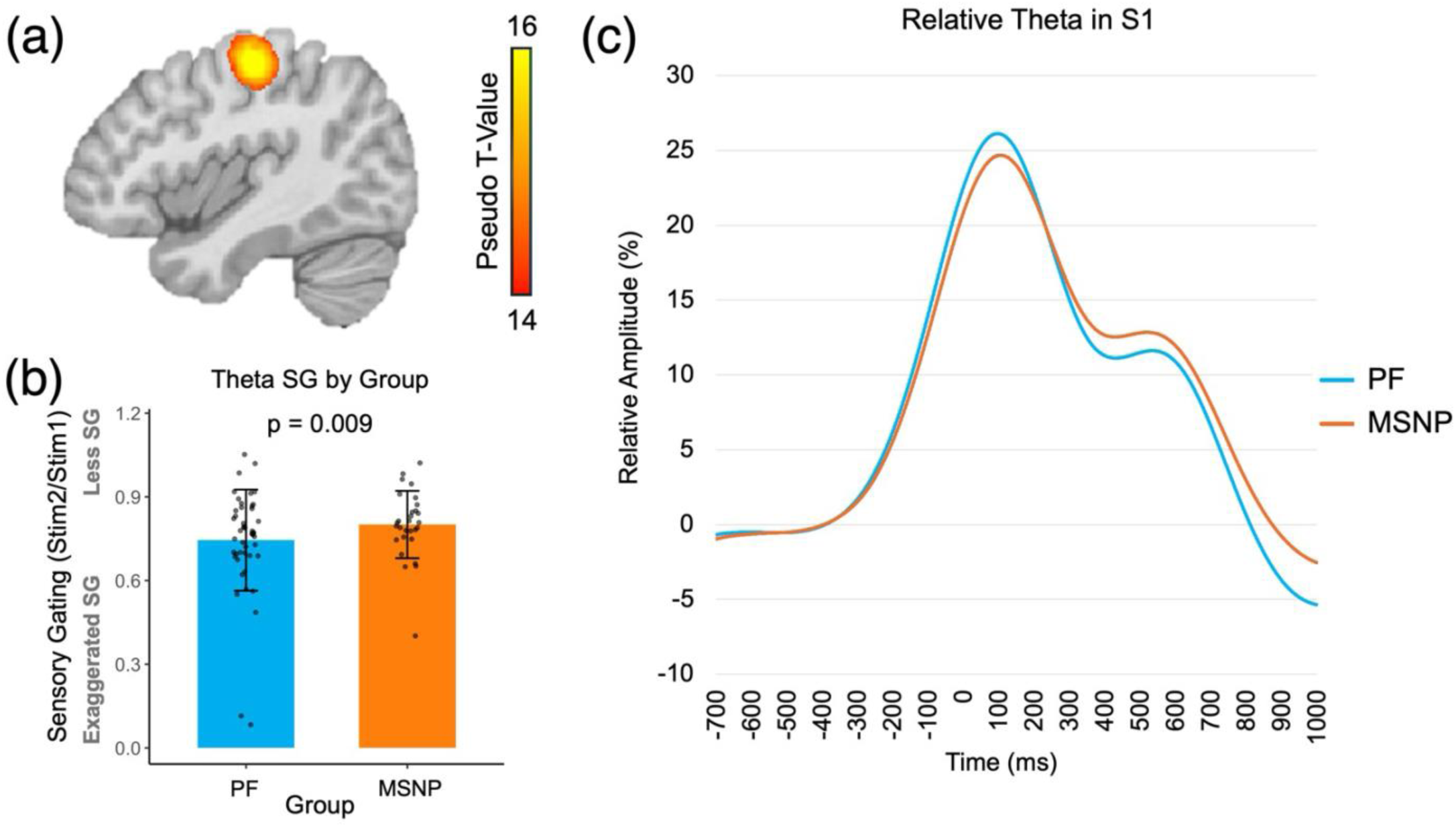
Group differences in theta SG. (a) The grand-averaged beamformer map (pseudo-*t*) collapsed across both stimulations and all participants in the theta band (4-7 Hz) revealed a cluster within contralateral S1. Time series (i.e., virtual sensor) data were extracted from the peak voxel in this cluster, per participant, and used to calculate the GR (Stim2/Stim1). (b) Theta SG was significantly reduced in adults with MSNP (orange) relative to PF adult controls (blue), above and beyond the effects of age and fibromyalgia (*p* = 0.009). Values of SG closer to 1 indicate less of a difference in theta response amplitude between Stim1 and Stim2 (i.e., less gating). Bars depict raw (not adjusted) means for PF controls and individuals with MSNP. Error bars represent ±1 standard deviation of the mean. (c) The relative amplitude time series of theta within contralateral S1 are plotted by group for visualization purposes. MSNP = moderate-to-severe nociceptive pain; PF = pain free.

### 3.3 Increased somatosensory gamma responses in individuals with MSNP

Given past literature suggesting heightened bottom-up processing of sensory stimuli in pain conditions (Apkarian et al., 2005; Kenefati et al., 2023; Ploner et al., 2017; Zebhauser et al., 2023), we evaluated whether individuals with MSNP exhibited heightened gamma responses to innocuous, repetitive stimulation. Gamma metrics (Stim1 relative amplitude, Stim2 relative amplitude, GR, and baseline absolute amplitude) were calculated from the peak voxel timeseries and entered into GLMs with group (MSNP, PF) as the variable of interest and age and fibromyalgia diagnosis as variables of no interest. A significant group effect was found on gamma Stim2 relative response amplitude above and beyond the effects of age and fibromyalgia (β = 0.523, t(3, 72) = 2.083, *p* = 0.041), such that individuals with MSNP had stronger gamma responses in contralateral S1 relative to PF individuals (Figure 3b). However, these group effects were not present when controlling for current pain intensity (i.e., day of scan self-report pain rating) (β = 0.388, t(4, 71) = 1.296, *p* = 0.199). No other gamma metrics were found to significantly differ by group and group-by-age interactions were not observed (all *p*’s > 0.05).

**Figure 3.**
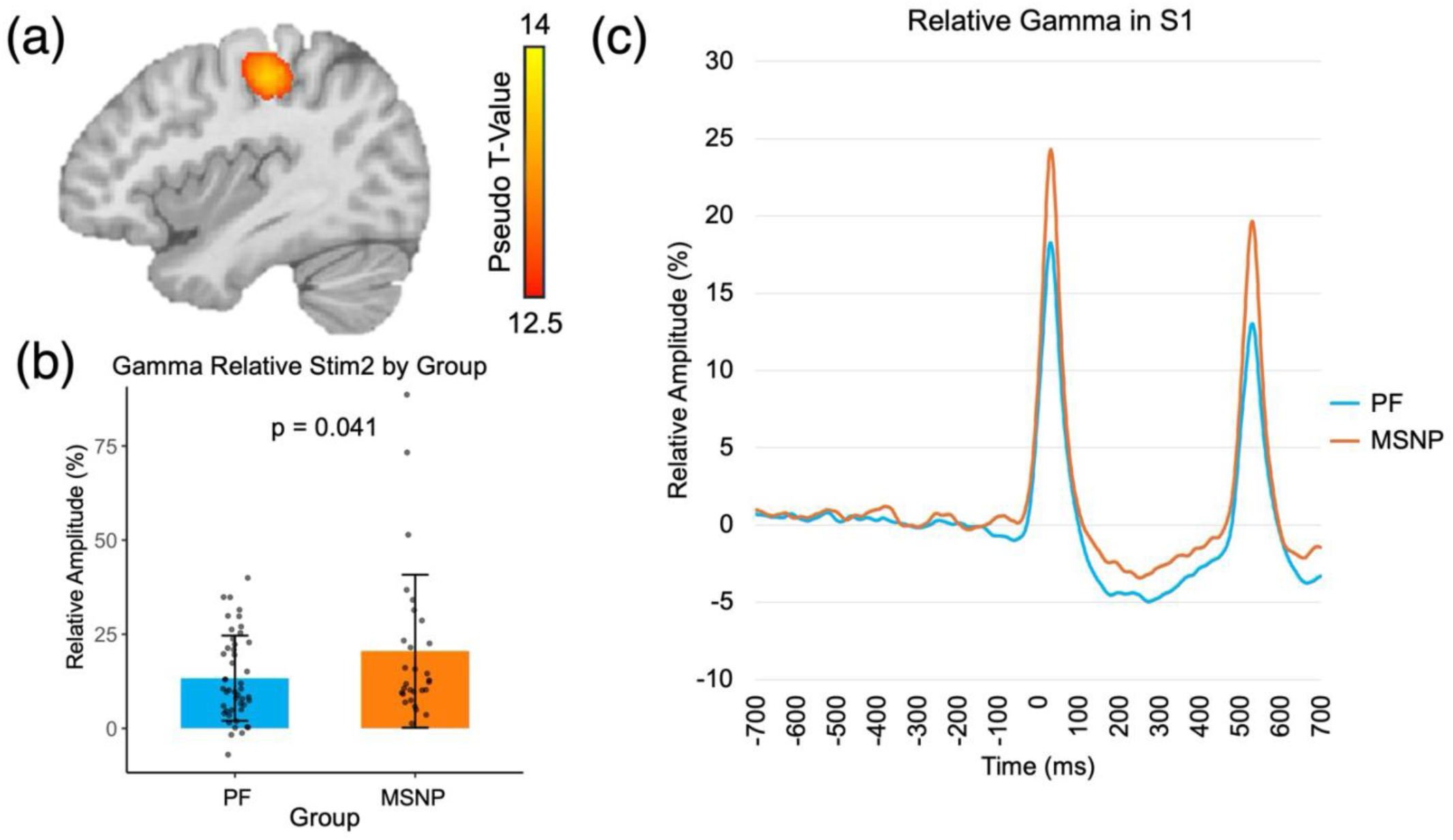
Group differences in Stim2 gamma response amplitude. (a) The grand-averaged beamformer map (pseudo-*t*) collapsed across both stimulations and all participants in the gamma band (30-90 Hz) revealed a cluster within contralateral S1. Time series (i.e., virtual sensor) data were extracted from the peak voxel, per participant, and used to obtain Stim1 and Stim2 relative response amplitudes. (b) Individuals with MSNP (orange) showed significantly stronger gamma responses in contralateral S1 following Stim2 relative to PF individuals (blue), above and beyond the effects of age and fibromyalgia (*p* = 0.041). Bars depict non-adjusted means for PF controls and individuals with MSNP. Error bars represent ±1 standard deviation of the mean. (c) The relative amplitude time series of gamma within contralateral S1 are plotted by group for visualization purposes. MSNP = moderate-to-severe nociceptive pain; PF = pain free.

### 3.4 Decreased somatosensory alpha in individuals with MSNP

Given past literature suggesting blunted top-down processing of sensory stimuli in pain conditions (Ahn et al., 2019; Kenefati et al., 2023; Ploner et al., 2017; Zebhauser et al., 2023), we evaluated whether individuals with MSNP exhibited reduced alpha responses to the SG paradigm. Similar to the other bands, the peak voxel timeseries were used to calculate gating and oscillatory metrics (Stim1 relative amplitude, Stim2 relative amplitude, GR, and baseline absolute amplitude) which were entered into GLMs with group (MSNP, PF) as the variable of interest and age and fibromyalgia diagnosis as variables of no interest. A significant group effect was found on alpha Stim1 relative response amplitude above and beyond the effects of age and fibromyalgia (β = 0.249, t(3, 72) = 2.018, *p* = 0.047) such that adults with MSNP demonstrated weaker responses relative to PF adults (Figure 4b,c). Note that the alpha response is an event-related desynchronization, so an increased relative amplitude is a value that is less negative and closer to zero, thus, resembling less alpha engagement from baseline. Additionally, alpha baseline activity was observed to have a significant group effect (β = -0.255, t(3, 72) = -2.110, *p* = 0.038), such that individuals with MSNP displayed weaker spontaneous alpha in contralateral S1 relative to PF individuals above and beyond the effects of age and fibromyalgia (Figure 4d,e). No other alpha metrics were found to significantly differ by group, and there were no group-by-age interactions (all *p*’s > 0.05).

**Figure 4.**
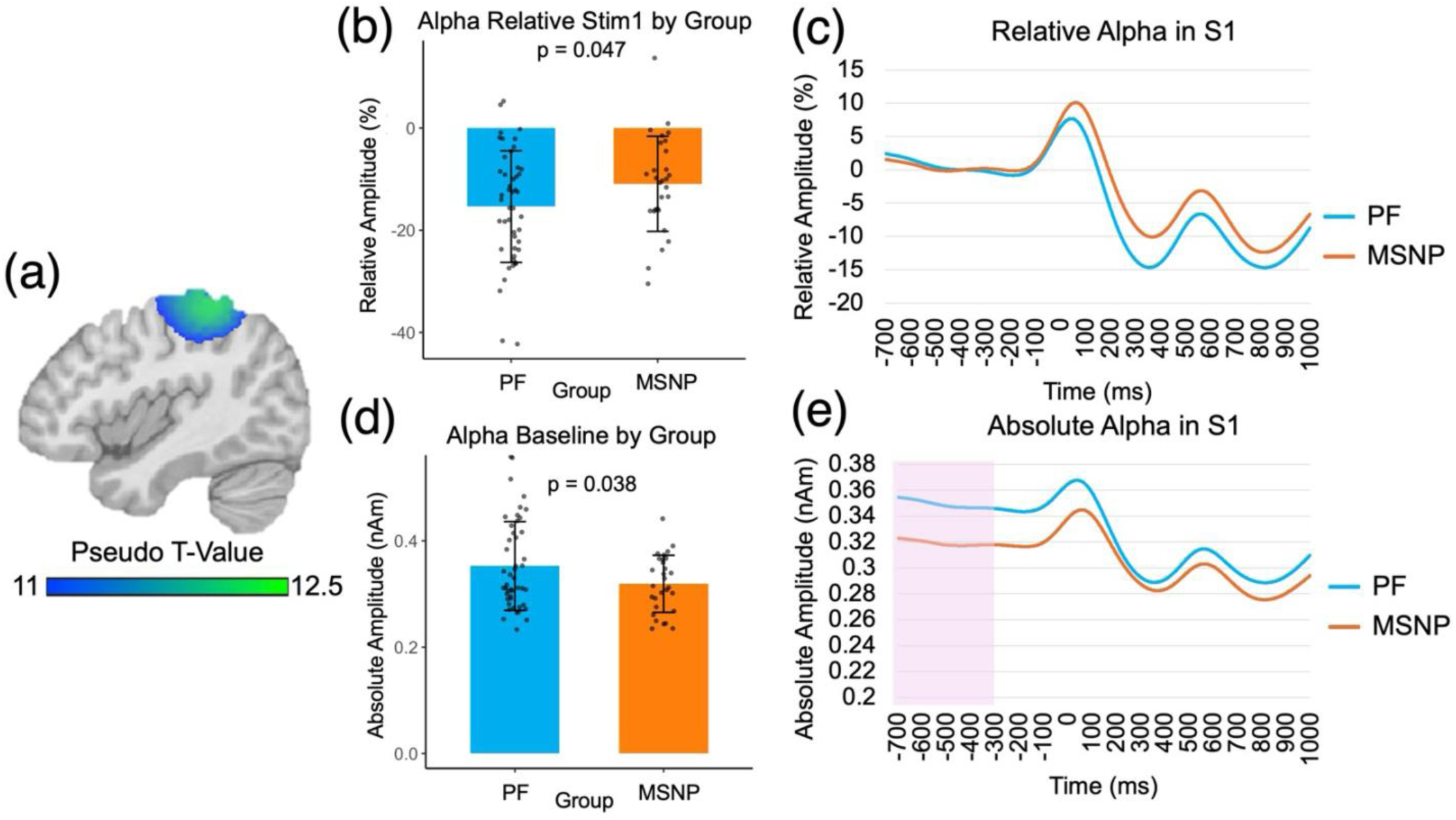
Group differences in alpha response amplitude and spontaneous alpha. (a) The grand-averaged beamformer map (pseudo-*t*) collapsed across both stimulations and all participants in the alpha band (8-13 Hz) revealed a cluster within contralateral S1. Time series (i.e., virtual sensor) data were extracted from the peak voxel, per participant. (b) Individuals with MSNP (orange) displayed significantly weaker alpha responses in contralateral S1 following Stim1 relative to PF individuals (blue), above and beyond the effects of age and fibromyalgia (*p* = 0.047). Bars depict means for raw (not adjusted) relative amplitude for PF controls and individuals with MSNP. Error bars represent ±1 standard deviation of the mean. Note that negative values reflect event-related desynchronizations, or decreases relative to baseline. (c) The relative amplitude time series of alpha within contralateral S1 are plotted by group for visualization purposes. (d) Adults with MSNP showed significantly weaker spontaneous alpha within contralateral S1 relative to PF adults above and beyond the effects of age and fibromyalgia (*p* = 0.038). Bars depict means of raw absolute amplitude for PF controls and individuals with MSNP. Error bars represent ±1 standard deviation of the mean. (e) Contralateral S1 alpha absolute amplitude time series are plotted by group with the baseline interval (−700 to -300 ms) shaded in pink. MSNP = moderate-to-severe nociceptive pain; PF = pain free.

### 3.5 Decreased spontaneous somatosensory alpha correlates with increased pain intensity in MSNP

Based off past literature suggesting sensory processing dysfunction and pain perception are correlated (Ahn et al., 2019), in the MSNP group only, we compared theta GR, Stim1 alpha response amplitude, and baseline alpha absolute amplitude to pain intensity ratings and corrected for multiple comparisons. A GLM revealed that spontaneous alpha within contralateral S1 was significantly and negatively correlated with pain intensity (β = -0.453, t(25) = -2.581, *p* = 0.016), above and beyond the effects of age and fibromyalgia (Figure 5b). That is, in individuals with MSNP, those who displayed weaker spontaneous somatosensory alpha tended to report more severe pain. Figure 5a displays the raw amplitude and categorical pain intensity ratings for visualization purposes, whereas Figure 5b displays the residual plot for the absolute amplitude and pain intensity. The other oscillatory metrics were not significantly correlated with pain intensity. Additionally, analyses did not yield significant group-by-age effects (*p*’s > 0.05).

**Figure 5.**
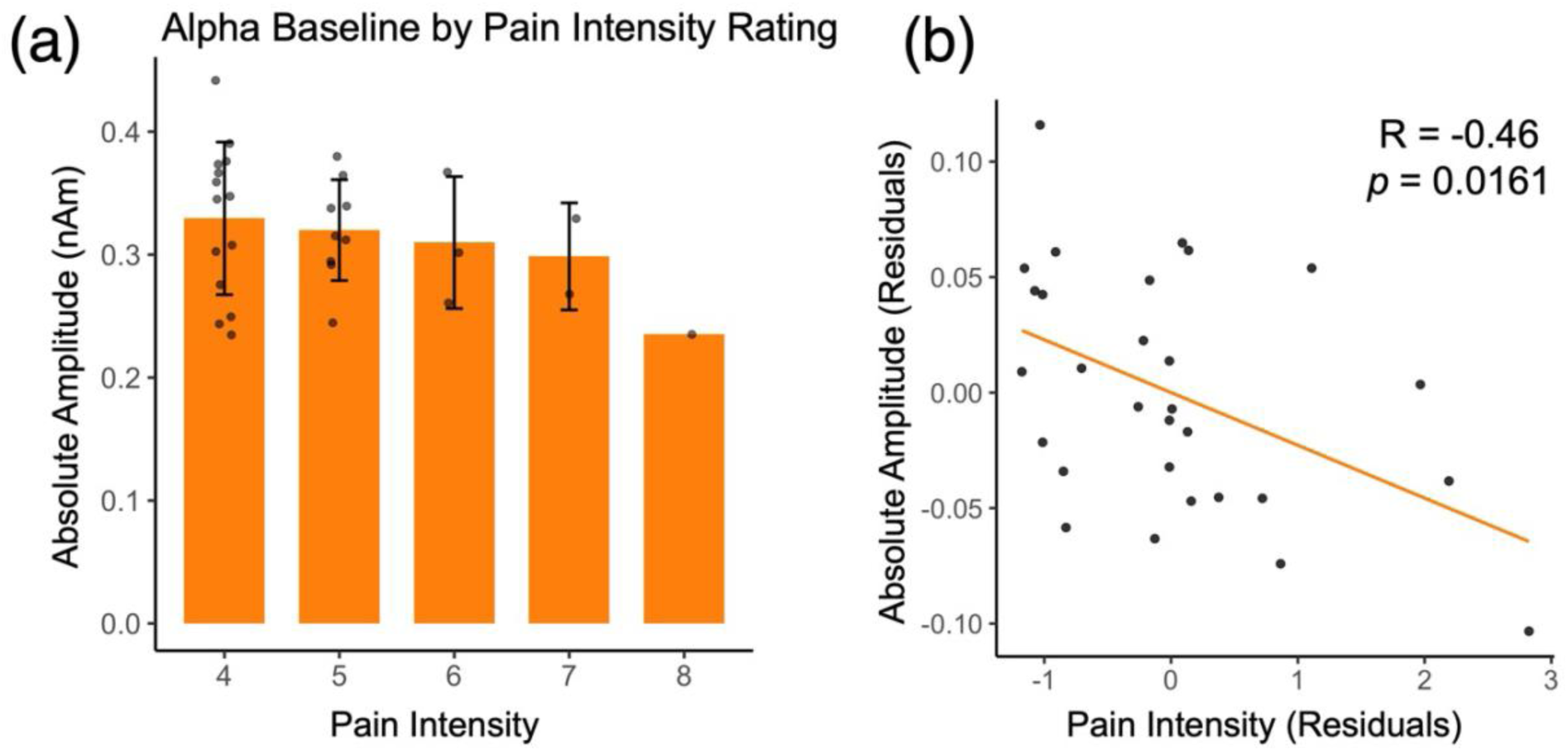
Spontaneous alpha within S1 correlates with pain intensity ratings in individuals with MSNP. (a) The bar plot visualizes spontaneous alpha (8-13 Hz) within contralateral S1 by pain intensity categories. The y-axis represents absolute amplitude of alpha during the baseline period (−700 to -300 ms). The x-axis reflects pain intensity by subjective rating measured by the PROMIS-29 instrument. Pain intensity of 8 was reported by only one participant; therefore, no variance bar is shown. Error bars represent ±1 standard deviation of the mean. (b) The scatterplot demonstrates the relationship between S1 alpha absolute amplitude (y-axis) and pain intensity (x-axis), accounting for age and fibromyalgia variance, in the MSNP group. A significant negative relationship was observed (*p* = 0.016). MSNP = moderate-to-severe nociceptive pain.

Lastly, MRI analyses did not identify significant structural-functional relationships (see Supplementary Material).

## 4 Discussion

This study is the first to identify that in individuals with MSNP, functional inhibition of redundant innocuous stimuli is reduced, bottom-up processing of innocuous stimuli is increased, and oscillatory mechanisms engaged in top-down attentional control are blunted and associated with pain intensity. Specifically, we investigated the impact of MSNP on SG, a functional inhibitory neural mechanism, in the contralateral (left) S1. We first identified that theta SG was significantly reduced (i.e., less functional inhibition) in individuals with MSNP compared to PF controls. We further demonstrated that gamma responses to innocuous stimulation were heightened, while alpha responses and spontaneous alpha were weakened in individuals with MSNP compared to PF controls. Lastly, spontaneous alpha was correlated with pain intensity ratings. These findings substantially expand our understanding of the pathophysiology of nociceptive pain conditions and further the idea that cortical disinhibition is linked to pain conditions.

Past studies have highlighted S1 as an important region for bottom-up and top-down mechanisms in both innocuous sensory processing and nociceptive pain modulation (Apkarian et al., 2005; Bushnell et al., 1999; Gross et al., 2007; Ploner et al., 2006). Further, S1 is known to exhibit SG, a functional inhibitory mechanism, in both time (i.e., neural evoked response) and time-frequency (i.e., neural oscillatory response) domains (Cromwell et al., 2008; Wiesman et al., 2017). Multiple studies have investigated S1 SG in the time-domain in either nociplastic or complex pain conditions (de Tommaso et al., 2011; Lenz et al., 2011; Lim et al., 2015; Montoya et al., 2006). Both of these pain conditions were found to have reduced SG, suggesting disinhibition of innocuous somatosensory stimuli, theorized to be due to central sensitization. In individuals with nociplastic pain, reduced SG was correlated with increased clinical pain symptomatology (Lim et al., 2015). This reduction of SG is thought to reflect thalamocortical disruptions, specifically thalamic disinhibition (Lim et al., 2015; Montoya et al., 2006). On the other hand, SG in the oscillatory domain has been tied to higher cognitive mechanisms beyond perception, such as attention (Wiesman et al., 2021; Wiesman & Wilson, 2020). In the present study, we identified that individuals with MSNP exhibited reduced SG of theta responses, but no group differences were observed in the gating of alpha, beta, or gamma responses. Specifically, theta SG is known to increase when individuals are paying attention to the somatosensory stimuli, opposed to a visual distractor task (Wiesman & Wilson, 2020). In our study, decreased theta SG in individuals with MSNP, compared to PF controls, may reflect altered cortical processing of somatosensory input. Theta oscillations during somatosensory stimulation are thought to play an important role in coordinating early sensory processing and regulating the integration of incoming sensory information (Beese et al., 2017; Fan et al., 2007; Fries, 2015; Gedankien et al., 2023; Gu et al., 2017; Iwamoto et al., 2021; Jensen, 2006; Kenefati et al., 2023; Lim et al., 2016; Nyhus & Curran, 2010; Summerfield & Mangels, 2005; Taesler & Rose, 2016; Tendler & Wagner, 2015; Wang et al., 2018; Wang, 2010; Wulff-Abramsson et al., 2025). In the somatosensory system, theta activity has been linked to the temporal organization of stimulus processing and the modulation of cortical responsiveness to repeated inputs (Kenefati et al., 2023; Wiesman & Wilson, 2020; Wulff-Abramsson et al., 2025). Reduced theta SG in individuals with MSNP may therefore indicate disrupted neural mechanisms supporting the efficient filtering or regulation of innocuous somatosensory signals within S1. Interestingly, gamma SG was not affected in individuals with MSNP compared to PF controls, suggesting that certain aspects of local cortical inhibitory processing may remain relatively preserved. Overall, these findings suggest that individuals with MSNP exhibit altered short-term neural adaptation to repeated non-nociceptive somatosensory stimulation, potentially reflecting changes in oscillatory dynamics that support sensory processing within S1. However, only one prior study has investigated theta SG (Wiesman & Wilson, 2020), and thus, future studies are needed to better evaluate the role of theta SG in S1 and further, in pain conditions.

In our current study, we identified exaggerated bottom-up processing of the second stimulus, indicated by increased gamma Stim2 response amplitude for individuals with MSNP compared to PF controls. Interestingly, past studies have implicated nociplastic and neuropathic pain conditions with exaggerated resting-state gamma activity, and these findings have also been used to suggest thalamocortical oscillation dysfunction (Lim et al., 2016; Zhou et al., 2018). However, to our best knowledge, no studies exist that investigate gamma in S1 in nociceptive pain conditions upon stimulus presentation with concurrent neuroimaging and peer-reviewed pain questionnaires. Thus, all previous studies investigating gamma oscillations and nociception depended on delivering acute or tonic noxious stimuli to healthy participants (Gross et al., 2007; Hauck et al., 2007; Li et al., 2016; Peng et al., 2014; Rossiter et al., 2013; Zhang et al., 2012). The tonic studies are thought to better capture pain conditions. In congruence with our findings, they have reported exaggerated gamma activity in S1 (Li et al., 2016; Peng et al., 2014) and additionally linked this increased activity to pain perception (Peng et al., 2014). Still other studies report increased gamma activity in prefrontal regions as key markers for pain perception (Schulz et al., 2015), lending to complicated interpretations of these high frequency oscillations in pain. However, and importantly, tonic pain paradigms exist on much shorter timescales (i.e., minutes) than MSNP (i.e., days) and are not accompanied by the same cognitive burden implications as an individual living with a nociceptive pain condition (e.g., arthritis). Our findings suggest that in individuals with MSNP, bottom-up processing of innocuous somatosensory stimuli is heightened, yet these responses are not associated with subjective pain intensity ratings. Notably, the observed group differences were no longer present when controlling for current pain intensity (i.e., pain reported on the day of the MEG scan). However, current pain state was not a significant predictor of group effects, nor did it mediate these effects. Future studies with a larger sample size would help elucidate whether collinearity between group and current pain state impacted the current interpretation. In contrast, group differences in alpha and theta activity were not affected by current pain state. Future studies should consider including current pain intensity as a covariate when examining oscillatory responses in pain populations, as failure to do so may contribute to inconsistent findings across the literature. Consistent with this possibility, an EEG study examining gamma activity in individuals with mixed chronic pain compared to healthy controls reported that gamma oscillations over central electrodes were associated with perceived pain intensity in healthy controls, but not in individuals with chronic pain (Bassez et al., 2020). These findings challenge the notion that gamma oscillations consistently track pain perception in chronic pain populations. Although the present study did not assess responses to noxious stimuli, our results similarly suggest that gamma responses to innocuous somatosensory stimulation do not explain variance in the severity of subjectively reported nociceptive pain.

Our current study identified that weaker spontaneous alpha within S1 was correlated with increased subjectively reported pain intensity, suggesting that alpha oscillations play an important role in pain disinhibition. Alpha oscillations in sensory regions are widely thought to reflect mechanisms that regulate the engagement and disengagement of behaviorally relevant sensory information, supporting top-down cognitive control over somatosensory processing (Fries, 2015; Haegens et al., 2011; Haegens et al., 2012; Jensen, 2024; Klimesch, 2012; Suffczynski et al., 2001; van Ede et al., 2014; van Kerkoerle et al., 2014; Wiesman & Wilson, 2020). For example, Brickwedde et al. observed increased absolute alpha power in S1 led to enhanced tactile perceptual learning efficiency; further, decreased absolute alpha power in S1 correlated with impeded tactile learning, suggesting perceptual disinhibition and inefficient somatosensory processing (Brickwedde et al., 2019). Notably, this relationship was specific to the alpha band and was not observed in theta, beta, or gamma frequencies (Brickwedde et al., 2019). Similarly, an EEG study investigating individuals with mixed pain found that spontaneous alpha band activity in centroparietal leads, near S1, correlated with clinical pain metrics (Ahn et al., 2019). In the same study, they confirmed increased alpha band activity in sensors near S1 via transcranial alternating current stimulation (bifrontal 10 Hz-tACS, corresponding to leads F3 and F4) and found that increased absolute alpha power was correlated with pain relief, identifying this modulation as a potential treatment outcome or marker (Ahn et al., 2019). In contrast, in healthy individuals, increasing alpha power was not linked to decreases in pain perception, suggesting that the proposed alpha-based cognitive control mechanism is not observed during acute nociceptive responses (i.e., transient, laser-induced pain) in healthy controls (who do not reflect ongoing pain states) (Hohn et al., 2025). Taken together, these findings suggest that individuals with MSNP exhibit less neural inhibitory mechanisms, and that this disinhibition profile reflects bottom-up and top-down somatosensory processing dysfunction.

This study has several limitations. First, because our design is cross-sectional, we cannot determine whether altered SG represents a cause or consequence of nociceptive pain. Longitudinal studies will be necessary to clarify the temporal relationship between disrupted inhibitory processing and the development or maintenance of pain. Second, the MSNP group included individuals with a range of clinical conditions (e.g., arthritis, surgical pain, etc.), which may introduce heterogeneity and contribute to variability in neural responses. However, the presence of group-level differences suggests that common central mechanisms may nonetheless underlie nociceptive pain across conditions, and the use of validated and reliable pain questionnaires supports the nociceptive pain category. Third, SG was assessed using innocuous stimulation, which does not directly engage nociceptive pathways and instead reflects more general somatosensory inhibitory processing. However, this approach is important to understand how cortical processing of MSNP impacts cortical processing of innocuous stimuli, since both share neural resources in S1. Finally, we did not perform quantitative sensory testing (QST) in the majority of participants. As a result, our characterization of pain relied primarily on self-reported measures of pain intensity, which limits our ability to objectively assess sensory function or determine whether participants exhibited altered peripheral nociceptive sensitivity. Future studies incorporating QST would help clarify how oscillatory dynamics and sensory gating relate to objective measures of somatosensory and nociceptive processing. Finally, structural and diffusion analyses did not yield significant results, likely due to limited power and sample size (discussed in Supplementary Material).

## 5 Conclusion

In summary, this study provides initial evidence that individuals with nociceptive pain exhibit altered oscillatory dynamics in S1, characterized by reduced functional inhibition, heightened bottom-up responses to innocuous stimulation, and diminished alpha oscillatory activity associated with pain symptoms. These findings suggest that nociceptive pain is accompanied by cortical disinhibition and altered somatosensory processing, extending mechanistic frameworks that have largely been applied to neuropathic and nociplastic pain to individuals with MSNP. By demonstrating central inhibitory alterations in MSNP, this work supports the possibility that nociceptive pain may share overlapping neural features with nociplastic pain phenotypes, highlighting the potential for shared central mechanisms across traditionally distinct pain classifications. Alternatively, these findings may suggest that nociplastic phenotypes include nociceptive components that contribute to altered sensory processing. Although the present study focused on S1, pain processing involves a broader network of cortical and subcortical regions (e.g., insula, anterior cingulate, thalamus) that may also contribute to altered sensory gating and somatosensory processing. Future work should therefore examine how network-level dynamics across pain-related brain regions contribute to inhibitory dysfunction in nociceptive pain.

## 6 Data availability

Requests for access to data may also be submitted to the Dallas Hearts Study (DHS) Publications Committee according to established study procedures.

## 7 Author Contributions

ALP and EMD contributed to the development, design, and validation of the MEG methodology. FFY contributed to the development and design of the MRI methodology. UEM and JZ contributed to the design of the pain questionnaire methodology. MV completed the formal MEG and MRI data analyses, visualization, investigation, and wrote the original draft of the manuscript. MV, LG, and SW contributed to MRI data preprocessing, and ALP, EMD, MV, NMB, LG, and SW assisted in data collection. YX, TP, FFY, AMS, JAM, and ALP contributed to the supervision of the data analyses. YX, NMB, TP, LG, SW, FFY, UEM, JZ, AMS, EMD, JAM, and ALP provided edits to the manuscript.

## 8 Funding Declaration

The Harry S. Moss Heart Trust funded the DHMS through UTSW. The funders did not play a role in study design, data collection and analysis, publication decision, or manuscript preparation.

## 9 Declaration of Competing Interests

The authors do not have any competing interests to disclose.

## 10 Acknowledgements

We would like to acknowledge Adriana Ohm for her assistance with DHMS piloting and data collection.

## Notes

### Competing Interest Statement

The authors have declared no competing interest.

