## Supplemental materials for "Reduced Somatosensory Oscillatory Dynamics and Inhibition in Moderate-to-Severe Nociceptive Pain"

Supplementary Material

Methodology for surface-based morphometry analyses

Beyond functional metrics, MRI studies have shown that pain can affect the neurostructural properties of the brain. For instance, in healthy individuals exposed to repetitive noxious stimulation over eight days, gray matter (GM) increased in S1 and other pain-related regions, indicating neurostructural plasticity after a relatively brief but ongoing pain experience (Rodriguez-Raecke et al., 2009; Teutsch et al., 2008). These structural changes were accompanied by alterations in brain function to reduce pain perception and thus, exhibit behavioral pain habituation (Bingel et al., 2008; Bingel et al., 2007; Rodriguez-Raecke et al., 2009; Teutsch et al., 2008). Motivated by these findings, we examined whether structural properties of S1 GM were associated with the functional MEG alterations observed in individuals with moderate-to-severe nociceptive pain (MSNP) relative to pain-free (PF) controls.

Structural measures were derived from high-resolution T1-weighted anatomical scans processed using the automated recon-all pipeline in FreeSurfer (version 7.1.1) (Fischl, 2012). This pipeline performs several preprocessing and reconstruction steps, including intensity normalization, skull removal, tissue classification, and reconstruction of the cortical surface. Cortical thickness was calculated at each surface vertex as the distance between the white matter boundary and the pial surface. Gray matter volume estimates were obtained from the anatomical segmentation generated during the reconstruction process. Vertex-level measurements were subsequently mapped to anatomical regions using the Desikan–Killiany–Tourville atlas, allowing extraction of region-of-interest (ROI) values for the left postcentral gyrus. Because regional GM volume is influenced by overall head size, intracranial volume was included as a covariate in statistical models involving volumetric measures.

ROI-level cortical thickness and GM volume values from the postcentral gyrus were entered into linear regression models to test for group differences, while controlling for nuisance variables. In the event of significant group effects, these structural measures were planned for additional analyses examining relationships with MEG-derived somatosensory metrics. However, unlike the clear group differences observed in MEG-based sensory gating measures, no significant differences were detected between groups in either cortical thickness or GM volume of the postcentral gyrus (all $p$'s > 0.05). This pattern suggests that the functional alterations observed in S1 are unlikely to reflect large-scale structural differences in this region. Nevertheless, the relatively modest MSNP sample size limits statistical power, and subtle structural variations may only become detectable in larger cohorts.

Methodology for diffusion analyses

An advanced approach for characterizing brain microstructure is multi-shell diffusion-weighted magnetic resonance imaging (dMRI), which allows modeling of the non-Gaussian diffusion of water molecules within neural tissue. Unlike conventional diffusion approaches, multi-shell dMRI enables estimation of distinct tissue compartments and therefore provides more detailed information about brain microstructure. One commonly used framework is neurite orientation dispersion and density imaging (NODDI), which estimates parameters such as the intracellular volume fraction (ICVF) (Zhang et al., 2012). Importantly, multi-shell acquisitions allow assessment of microstructural properties not only within white matter but also in GM regions (Fukutomi et al., 2018; Nazeri et al., 2015; Zhang et al., 2012). Despite these advantages, diffusion-based investigations of pain have largely relied on diffusion tensor imaging (DTI) measures such as fractional anisotropy (FA) (Lieberman et al., 2014; Torrecillas-Martinez et al., 2020). While widely used, DTI assumes Gaussian diffusion and therefore provides a limited representation of tissue complexity, particularly in regions containing crossing fibers or smaller axons and dendritic structures (Fukutomi et al., 2018; Zhang et al., 2012). To address these limitations, the present study examined both conventional DTI metrics (FA and mean diffusivity [MD]) and NODDI-derived parameters (orientation dispersion [OD], isotropic volume fraction [ISOVF], and ICVF). These metrics were analyzed in relation to MEG-derived functional measures.

White matter microstructure was examined using Tract-Based Spatial Statistics implemented in FSL (Jenkinson et al., 2012; Smith et al., 2006). For each participant, FA and MD images were nonlinearly registered to the FMRIB58_FA template in standard space and resampled to 1 mm isotropic resolution. A group-average FA image was then generated and thinned to produce a white matter skeleton representing the centers of major fiber pathways (FA threshold > 0.2). Individual DTI maps were subsequently projected onto this skeleton for voxel-wise analysis.

NODDI parameters were estimated across the whole brain from the multi-shell diffusion data using custom MATLAB-based processing pipelines (Zhang et al., 2012). The resulting parametric maps were co-registered to each participant's T1-weighted anatomical image prior to statistical analysis.

Voxel-wise statistical testing was performed using permutation-based nonparametric inference (randomise; 5,000 permutations). Group differences were modeled while controlling for age and fibromyalgia diagnosis as nuisance covariates. An additional model was used to test potential group-by-age interaction effects. Statistical maps were enhanced using threshold-free cluster enhancement, and results were corrected for multiple comparisons using family-wise error correction with a significance threshold of $p < 0.05$. No significant group differences or associations were observed across any diffusion-derived metrics following correction for multiple comparisons (all $p$'s > 0.05). However, given the modest sample size of the current cohort, future studies employing larger samples will be necessary to determine whether subtle microstructural alterations may be detectable using multimodal imaging approaches.
